# Thrombocyte inhibition restores protective immunity to mycobacterial infection in zebrafish

**DOI:** 10.1101/338111

**Authors:** Elinor Hortle, Khelsey E. Johnson, Matt D. Johansen, Tuong Nguyen, Jordan A. Shavit, Warwick J. Britton, David M. Tobin, Stefan H. Oehlers

## Abstract

Infection-induced thrombocytosis is a clinically important complication of tuberculosis (TB). Recent studies have separately highlighted a correlation of platelet activation with TB severity and utility of aspirin as a host-directed therapy for TB that modulates the inflammatory response. Here we investigate the possibility that the beneficial effects of aspirin are related to an anti-platelet mode of action. We utilize the zebrafish-*Mycobacterium marinum* model to show mycobacteria drive host hemostasis through the formation of granulomas. Treatment of infected zebrafish with aspirin or platelet-specific glycoprotein IIb/IIIa inhibitors reduced mycobacterial burden demonstrating a detrimental role for infection-induced thrombocyte activation. We found platelet inhibition reduced thrombocyte-macrophage interactions and restored indices of macrophage-mediated immunity to mycobacterial infection. Pathological thrombocyte activation and granuloma formation were found to be intrinsically linked illustrating a bidirectional relationship between host hemostasis and TB pathogenesis. Our study illuminates platelet activation as an efficacious target of anti-platelets drugs including aspirin, a widely available and affordable host-directed therapy candidate for tuberculosis.

**Key Points:**

1. Inhibition of thrombocyte activation improves control of mycobacterial infection.
2. Inhibition of thrombocyte activation reduces thrombocyte-macrophage interactions and improves indices of macrophage immune function against mycobacterial infection.

## Introduction

*Mycobacterium tuberculosis* is the world’s most lethal pathogen, causing nearly 2 million deaths each year ^1^. The increasing incidence of both multi- and extremely-drug resistant tuberculosis (TB) urgently require the development of therapeutics that overcome the shortcomings of conventional antibiotics. Pathogenic mycobacteria co-opt numerous host pathways to establish persistent infection, and subversion of these interactions with host-directed therapies (HDTs) has been shown to reduce the severity of infection in animal models. For example, we have recently shown that mycobacteria induce host angiogenesis and increase host vascular permeability; blockade of either of these processes reduced both the growth and spread of bacteria ^2,3^. Therefore, host processes co-opted by the bacteria provide attractive targets for novel TB treatments. One such pathway may be hemostasis.

Thrombocytosis has long been recognized as a biomarker for advanced TB, and infection is often accompanied by the induction of a hyper-coagulable state, resulting in increased risk of deep vein thrombosis and stroke ^4,5^. Recent evidence hints that mycobacteria may drive this process, and that it may aid their growth. For example, cell wall components from *M*. *tuberculosis* can induce expression of tissue factor - an important activator of coagulation - in macrophages ^6^. In mice and humans markers of platelet activation are upregulated during *M*. *tuberculosis* infection ^7,8^, and it has been shown *in vitro* that interaction with activated platelets increases the conversion of infected macrophages into cells permissive for bacterial growth ^7,9^. To date the pathogenic roles of hemostasis have not been studied in an intact *in vivo* model of mycobacterial infection.

Here we used the zebrafish-*M*. *marinum* model to investigate the role of host thrombocytes in mycobacterial infection. We present evidence that while coagulation and thrombocyte activation are both driven by mycobacteria, it is only infection-induced activation of thrombocytes that specifically compromises protective immunity through direct thrombocyte-macrophage interactions.

## Methods

### Zebrafish husbandry

Adult zebrafish were housed at the Garvan Institute of Medical Research Biological Testing Facility (St Vincent’s Hospital AWC Approval 1511) and embryos were produced by natural spawning for infection experiments at the Centenary Institute (Sydney Local Health District AWC Approval 2016-022). Zebrafish embryos were obtained by natural spawning and embryos were raised at 28°C in E3 media.

### Zebrafish lines

Wild type zebrafish are the TAB background. Transgenic lines are: *Tg*(*fabp10a:fgb-EGFP)*^*mi4001*^ referred to as *Tg*(*fabp10a:fgb-EGFP)*^*10*^, *Tg*(*-6*.*0itga2b:eGFP)*^*la2*^ referred to as *Tg*(*cd41:EGFP)*^*11*^, *Tg*(*mfap4:tdTomato)*^*xt12*^ referred to as *Tg*(*mfap4:tdTomato)*^*12*^, Mutant allele *fga*^*mi*^ contains a 26 bp insertion in the *fibrinogen alpha chain* gene (manuscript in preparation).

### Infection of zebrafish embryos

Aliquots of single cell suspensions of midlog-phase *Mycobacterium marinum* M strain and ΔESX1 *M*. *marinum* were frozen at −80°C for use in infection experiments. Bacterial aliquots were thawed and diluted with phenol red dye (0.5% w/v). 10-15 nL was injected into the caudal vein or trunk of M-222 (tricaine)-anaesthetized 30-48 hpf embryos resulting in a standard infectious dose ∼400 fluorescent *M*. *marinum*. Embryos were recovered into E3 supplemented with 0.036 g/L PTU, housed at 28 °C and imaged on day 5 of infection unless otherwise stated.

### Drug treatments

Embryos were treated with vehicle control (DMSO or water as appropriate), 10 µg/ml aspirin, 20 µg/ml tirofiban, 10 µM eptifibatide, or 5 µM warfarin. Drugs and E3 were replaced on days 0, 2, and 4 days post infection (DPI) unless otherwise stated.

### Tail wound thrombosis assay

Three day post fertilization (DPF) embryos were treated over-night with anti-platelet drugs. They were anaesthetized, and then a small amount of their tail was removed with a scalpel. Embryos were imaged 4 hours post wounding and the number of GFP positive cells within 100 μm of the cut site was counted.

### Imaging

Live zebrafish embryos were anaesthetized in M-222 (Tricaine) and mounted in 3% methylcellulose for static imaging on a Leica M205FA or DM6000B fluorescence stereomicroscope. Image analysis was carried out with Image J Software Version 1.51j using fluorescent pixel counts and intensity measurements as previously described ^13^.

Video and timelapse imaging was carried out on anaesthetized embryos mounted in 0.75% low melting point agarose on a Leica M205FA or Deltavision Elite fluorescence microscope. Video editing was carried out with Image J Software Version 1.51j and iMovie.

### Axenic culture

A midlog culture of fluorescent *M*. *marinum* was diluted 1:100 and aliquoted into 96 well plates for drug treatment. Cultures were maintained at 28°C in a static incubator and bacterial fluorescence was measured in a plate reader.

### Morpholinos

Embryos were injected at the single cell stage with 1 pmol cMPL (5′-CAGAACTCTCACCCTTCAATTATAT-3′), or control morpholino (5′-CCTCTTACCTCAGTTACAATTTATA-3′).

### Clodronate liposome injections

Larvae were injected at 3 DPI (4 DPF) with 10 nl of 5 mg/ml clodronate liposomes or 5 mg/ml PBS vehicle liposomes by caudal vein injection.

### Oil-red O

Oil Red O lipid staining on whole mount embryos was performed and analyzed as previously described ^14,15^. Briefly, embryos were individually imaged for bacterial distribution by fluorescent microscopy, fixed, and stained in Oil Red O (0.5% w/v in propylene glycol). Oil Red O density was calculated by using the ‘measure’ function in Image J, and subtracting the mean brightness of a representative region within each granuloma from the mean brightness of a representative adjacent ‘background’ region.

### Statistics

All t-tests were unpaired t-tests with Welch’s correction. All ANOVA were ordinary one-way ANOVA, comparing the mean of each group with the mean of every other group, using Turkey’s multiple comparisons test with a single pooled variance. In cases where data was pooled from multiple experiments, data from each was normalized to its own within-experiment control (usually ‘DMSO’) before pooling. Outliers were removed using ROUT, with Q=1%.

## Results

### *Mycobacterium marinum* infection induces thrombocytosis in zebrafish

To determine if *M*. *marinum* induces thrombocytosis in zebrafish, we infected *Tg*)*cd41:GFP)* embryos, where thrombocytes are marked by GFP expression, with red fluorescent *M*. *marinum*. Infected embryos had significantly increased density of thrombocytes around the tail venous plexus where granulomas preferentially form (Figures 1A-B). Amongst infected embryos, there was a strong positive correlation between thrombocyte density and mycobacterial burden (Figure 1C).

**Figure 1:**
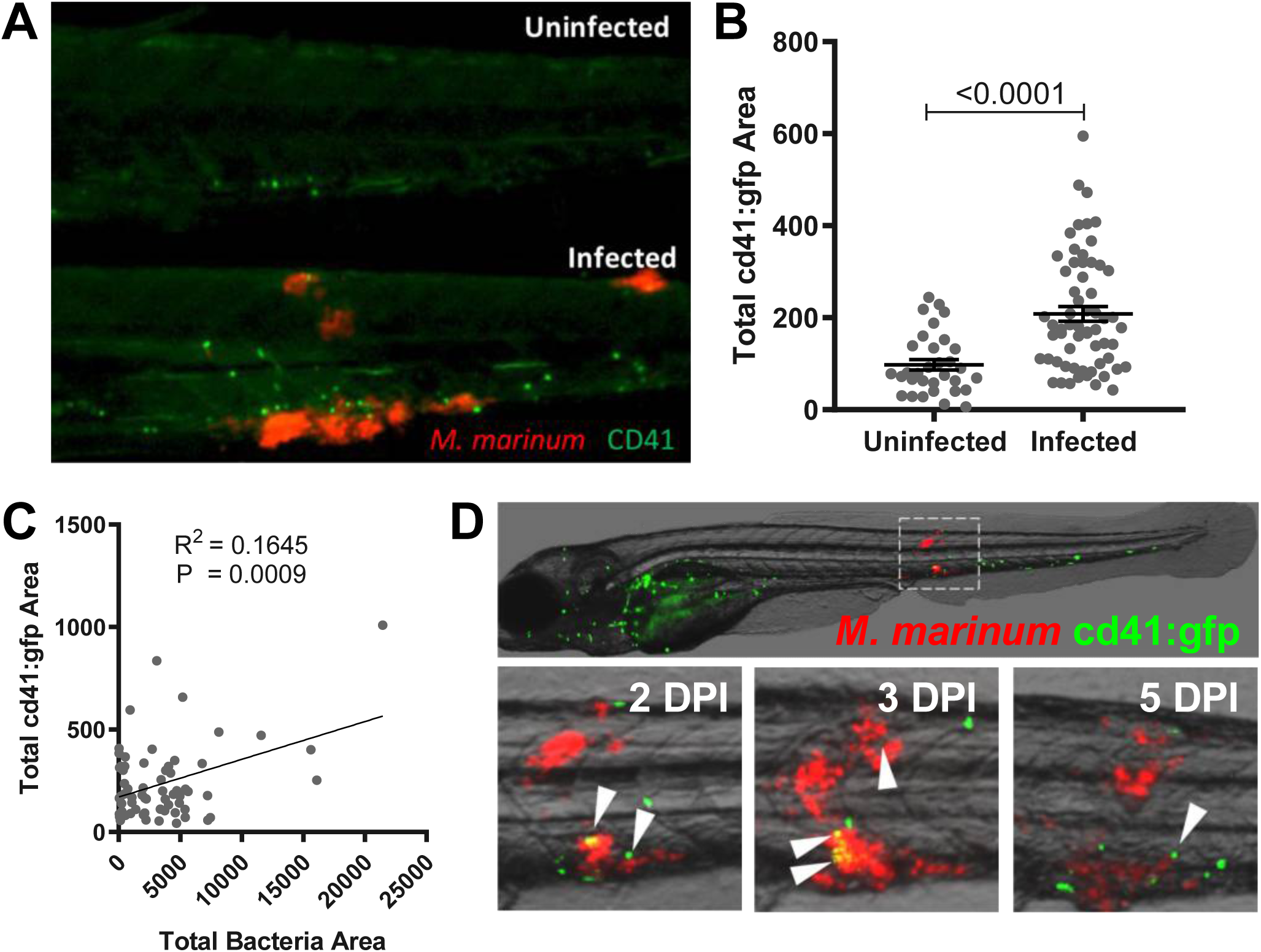
*Mycobacterium marinum* induces coagulation and thrombosis around sites of infection in zebrafish. A) Representative images of 5 DPI *Tg*(*cd41:GFP)* embryos infected with *M*. *marinum*-tdTomato, showing thrombocyte accumulation (green) at sites of bacterial infection (red). B) Quantification of total thrombocyte *Tg*(*cd41:GFP)* area within the tail of uninfected or WT *M*. *marinum*-tdTomato infected embryos. Graph shows mean ± SEM. Statistical analysis performed by T tests. C) Correlation between *M*. *marinum* bacterial burden and total thrombocyte *Tg*(*cd41:GFP)* area within the tail of infected embryos. P and R^2^ calculated using linear regression. D) Representative 2 DPI, 3 DPI and 4 DPI images of *Tg*)*cd41:GFP)* fish infected with *M*. *marinum*-tdTomato at 3 DPF. White arrowheads show thrombocyte (green) association with areas of bacterial growth (red). Thrombocytes not indicated with an arrowhead were circulating and not considered to be associated with bacteria.

Because the *cd41* promoter is active in non-motile thrombocyte precursors within their caudal hematopoietic tissue, we could not conclusively determine if these thrombocytes had actively migrated to and been retained at the site of infection ^11^. To determine if zebrafish thrombocytes are recruited to sites of mycobacterial infection, we performed trunk injection of *M*. *marinum* in *Tg*(*cd41:GFP)* at 3 days post fertilization (DPF), a time point after which mature thrombocytes are in the circulation. Embryos were then imaged at 2, 3, and 4 DPI. Rather than forming a stable and growing clot over a period of days, thrombocytes appeared to form transient associations with sites of infection, and new thrombocytes seemed to be retained at sites of infection in different locations each day (Figure 1D). We therefore recorded videos of *Tg*(*cd41:GFP)* embryos infected with *M*. *marinum*-tomato using a long pass GFP filter to capture bacterial and thrombocyte fluorescence simultaneously. Thrombocytes were most often observed on the edges of granulomas consistent with the location of granuloma-defining macrophages ^16^. They also formed short associations with sites of infection, sometimes lasting only 5-10 minutes (Supplementary Video 1). Therefore thrombocyte-granuloma interactions appear to be a conserved feature of mycobacterial infection across species.

### Anti-platelet drugs reduce mycobacterial burden in zebrafish

It has previously been reported that aspirin has a host-protective effect during TB infection ^17-20^. Most of these studies have focused on the fact that aspirin is a broadly acting nonsteroidal anti-inflammatory drug (NSAID) that is known to modulate infection-relevant prostaglandin metabolism ^21^. However, aspirin is also a widely-used platelet inhibitor, and we theorized this capacity may also play a role in the drug’s effectiveness against TB.

To test this hypothesis we first confirmed that aspirin’s protective effect was seen across species, by treating *M*. *marinum*-infected fish with aspirin by immersion. Mycobacterial burden was reduced by approximately 50% in aspirin-treated embryos (Figure 2A).

**Figure 2:**
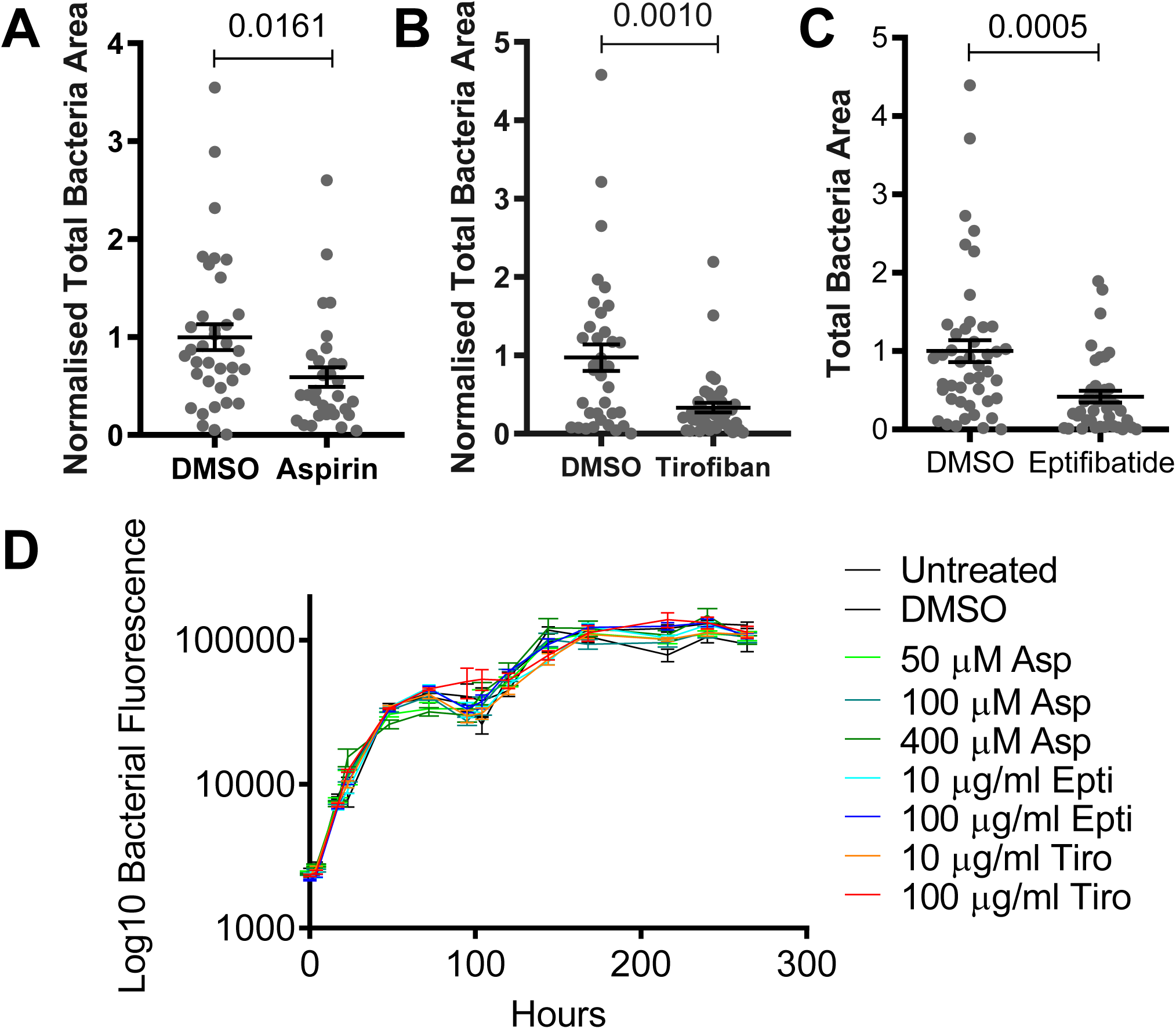
Anti-platelet drugs reduce bacterial burden in *M*. *marinum* infection. A) Quantification of bacterial burden in embryos treated with aspirin normalized to DMSO control. Data are combined results of two independent experiments. B) Quantification of bacterial burden in embryos treated with tirofiban normalized to DMSO control. Data are combined results of two independent experiments. C) Quantification of bacterial burden in embryos treated with eptifibatide or DMSO control. Data are combined results of two independent experiments. D) Quantification of bacterial growth by relative fluorescence in 7H9 broth culture supplemented with drugs as indicated. All graphs show Mean ± SEM. Statistical analyses performed by T tests.

To determine if the anti-platelet effects of aspirin treatment contribute to the reduced mycobacterial burden, we treated *M*. *marinum*-infected fish with the platelet-specific, small molecule glycoprotein IIb/IIIa inhibitors, tirofiban or eptifibatide. These drugs do not inhibit platelet activation and de-granulation, but rather inhibit activated platelets from binding to one-another, and to monocytes, via fibrinogen. Treatment with either glycoprotein IIb/IIIa inhibitor phenocopied aspirin by reducing bacterial burden providing direct evidence of a pathological role for thrombocyte activation in the immune response to mycobacterial infection (Figure 2B-C).

We next examined the cellular target of anti-platelet drugs in our infection system. We performed antibacterial testing of the anti-platelet drugs in axenic cultures of *M*. *marinum* and did not observe any effect on bacterial growth *in vitro* demonstrating host-dependent activity (Figure 2D).

### The anti-bacterial effect of anti-platelet drugs is thrombocyte dependent

We next confirmed that anti-platelet drugs inhibit thrombocytes in zebrafish using a tail wound thrombosis assay (Figure S1A). Anti-platelet drugs reduced the number of thrombocytes recruited to tail wound clots demonstrating conservation of their cellular target in zebrafish embryos (Figure S1B-C).

To determine if zebrafish thrombocytes are the conserved target for anti-platelet drugs in the zebrafish-*M*. *marinum* infection model, we inhibited thrombopoiesis by injection with a morpholino against the thrombopoietin receptor *cmpl* ^11^. Inhibition of thrombopoiesis did not affect the outcome of infection. However, both aspirin and tirofiban treatment failed to reduce bacterial burden in thrombocyte-depleted embryos, demonstrating thrombocytes are the cellular target of this drug in the zebrafish infection model (Figure 3A-B).

**Figure 3:**
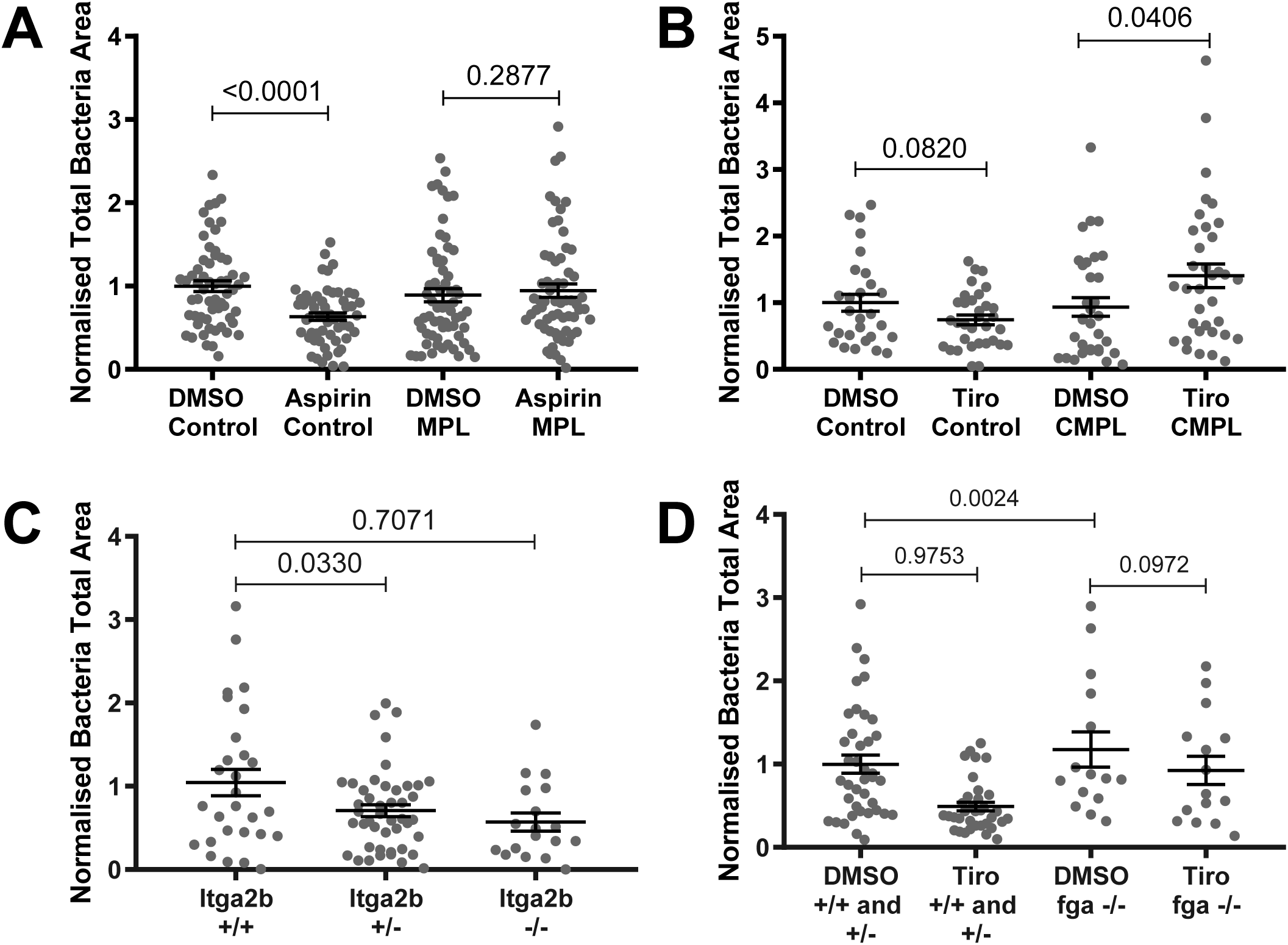
The anti-bacterial effect of anti-platelet drugs is thrombocyte dependent. A) Total fluorescence area of *M*. *marinum* bacteria in larvae injected with either control or cmpl morpholino (MO) to deplete thrombocytes, and then treated with aspirin. Values are normalized to DMSO-treated control MO larvae. Graphs show the combined results of 2 independent experiments. B) Total fluorescence area of *M*. *marinum* bacteria in larvae injected with either control or cmpl morpholino (MO) to deplete thrombocytes, and then treated with tirofiban (Tiro). Values are normalized to DMSO-treated control MO larvae. Graphs show the combined results of 2 independent experiments. C) Quantification of bacterial burden in in *itga2b* mutant embryos normalized to WT control. Data are combined results of 3 independent experiments. D) Quantification of bacterial burden in *fga*^-/-^ embryos treated with tirofiban, normalized to *fga*^*+/+*^ and *fga*^*+/-*^controls. Data are combined results of 2 independent experiments. All graphs show Mean ± SEM. Statistical analyses performed by ANOVA.

Eptifibatide and tirofiban were designed as specific inhibitors to prevent binding between Glycoprotein IIb/IIIa and fibrinogen in mammals ^22^, but nothing is known about their potential off-target effects in the zebrafish. Therefore, to confirm that disruption of Glycoprotein IIb/IIIa binding alone can reduce bacterial burden, we performed infection experiments in Glycoprotein IIb/IIIa knock-out (KO) transgenic embryos (*itga2b* mutants). The *itga2b*^*sa10134*^ allele caused a dose-dependent reduction in thrombocytes recruited to tail wound clots (Figure S1D), and KO of *itga2b* significantly reduced bacterial burden (Figure 3C).

Similarly, when we when we addressed the same question using a *fibrinogen alpha chain* (fga) mutant zebrafish line that does not produce mature fibrinogen, we saw that KO fish had significantly reduced bacterial burden (Figure 3D). Furthermore, while tirofiban was able to reduce bacterial burden in fga sufficient mutants, it did not reduce bacterial burden in fga KO. Together these data suggest that tirofiban is reducing bacterial burden by inhibiting binding between glycoprotein IIb/IIIa and fibrinogen.

### *Mycobacterium marinum* induces coagulation in zebrafish but inhibition of coagulation does not affect the outcome of infection

Seeing that there was a marginal decrease in bacterial burden in *fga-/-* embryos compared to WT and heterozygous clutchmates, we next sought to determine if *M*. *marinum* induces coagulation in zebrafish and the role of coagulation in *M*. *marinum* infection. We infected *Tg*(*fabp10a:fgb-EGFP)* transgenic embryos expressing EGFP-tagged fibrinogen beta (FGB) with *M*. *marinum*-tdtomato, and imaged the developing infection every 15 minutes from 3 days post infection (DPI), until 6 DPI (Supplementary Video 2). We observed that clots formed only at areas of bacterial growth, and that the size of the clots increased as the number of bacteria increased over the course of the infection (Figure 4A). When we infected fish with ΔESX1 mutant *M*. *marinum*, which lacks the ability to export key virulence proteins and does not form granulomas, we observed significantly reduced clot formation (Figures 4B).

**Figure 4:**
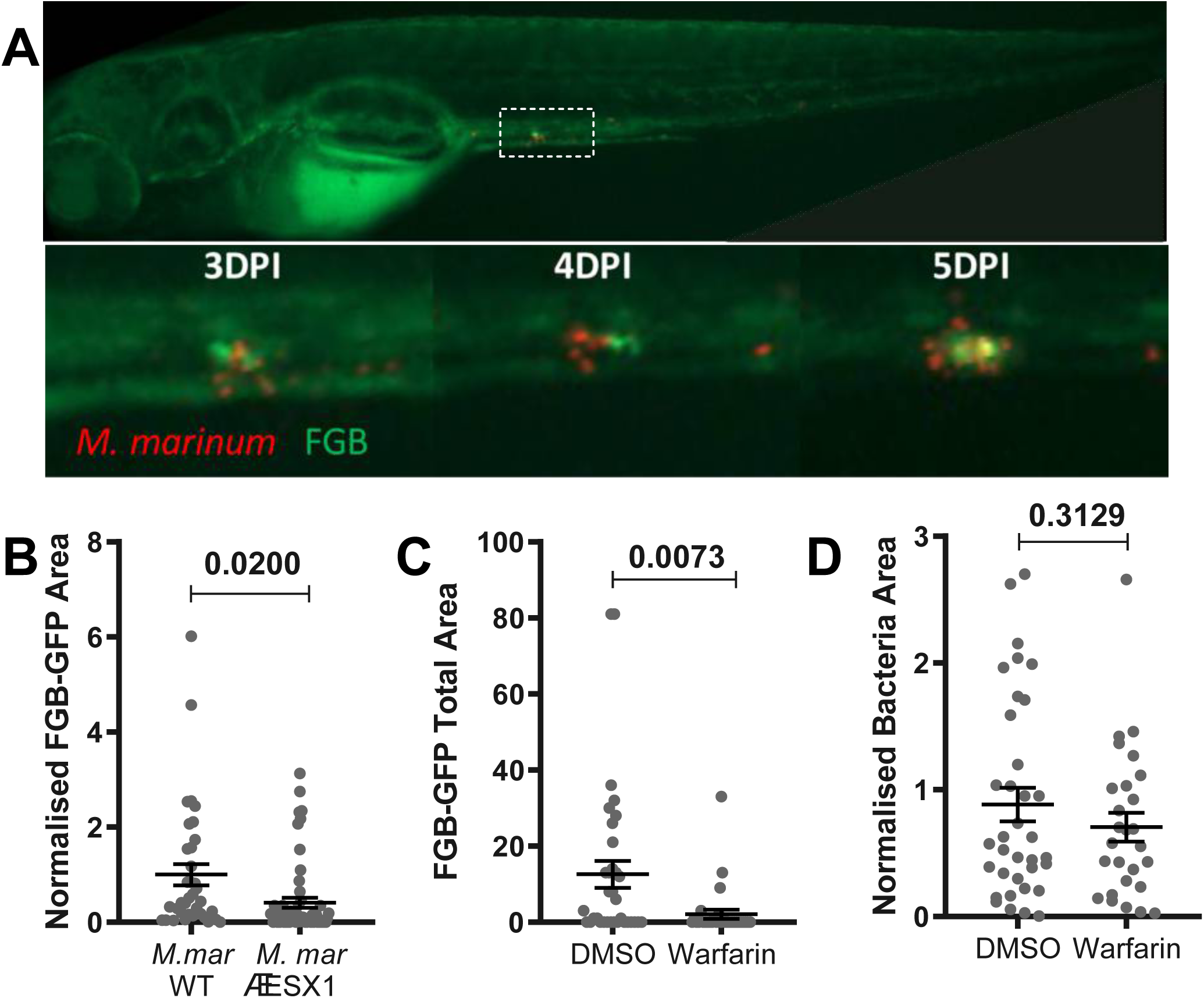
*Mycobacterium marinum* induces coagulation in zebrafish but inhibition of coagulation does not affect the outcome of infection. A) Representative images of a *Tg*(*fabp10a:fgb-EGFP)* embryo infected with *M*. *marinum*-tdTomato by caudal vein injection, showing clot formation (green) at sites of infection (red) at 3, 4, and 5 DPI. B) Quantification of clot formation in burden-matched ΔESX1 mutant-infected *Tg*(*fabp10a:fgb-EGFP)* embryos normalized to WT *M*. *marinum* control. Data show combined results from two independent experiments. C) Quantification of clotting in warfarin-treated *Tg*(*fabp10a:fgb-EGFP)* embryos. D) Quantification of bacterial burden in warfarin treated embryos, normalized to DMSO control. Data show combined results of two independent experiments. All graphs show mean ± SEM, statistical analyses by T tests.

These data suggested clotting could be driven by mycobacterial and may thus contribute to mycobacterial pathogenesis. To test this hypothesis, we treated infected embryos with the anti-coagulant warfarin to prevent clot formation. We did not observe significant changes in bacterial burden, suggesting coagulation itself does not affect bacterial growth within the host (Figures 4C-D). Together, these data demonstrate that while coagulation is a consequence of infection driven by a conserved mycobacterial pathogenicity program across host species, the effects of coagulation on mycobacterial pathogenesis vary between host species ^6^.

### Thrombocytes increase mycobacterial burden independently of coagulation

To assess the contribution of thrombocytes to infection-induced coagulation, we analyzed the formation of FGB-GFP clots in tirofiban-treated *Tg*(*fabp10a:fgb-EGFP)* embryos. Tirofiban visibly reduced total clot formation (Figure 5A). However, correction for relative bacterial burden suggested that the reduced clot formation was burden-dependent and thrombus formation was not additionally impacted by tirofiban treatment (Figure 5B). Therefore, we hypothesized that tirofiban was reducing bacterial burden independently of infection-induced coagulation. To investigate this hypothesis, we again used warfarin, which prevented clot formation during infection and did not affect bacterial burden (Figure 4C-D and 5C). As expected, the addition of warfarin to our tirofiban treatment model had no effect on the ability of tirofiban to reduce bacterial burden (Figure 5D), indicating that tirofiban acts through an independent process. This suggests that the protective effect of tirofiban occurs independently of fibrin clot formation, but requires the presence of soluble fibrinogen.

**Figure 5:**
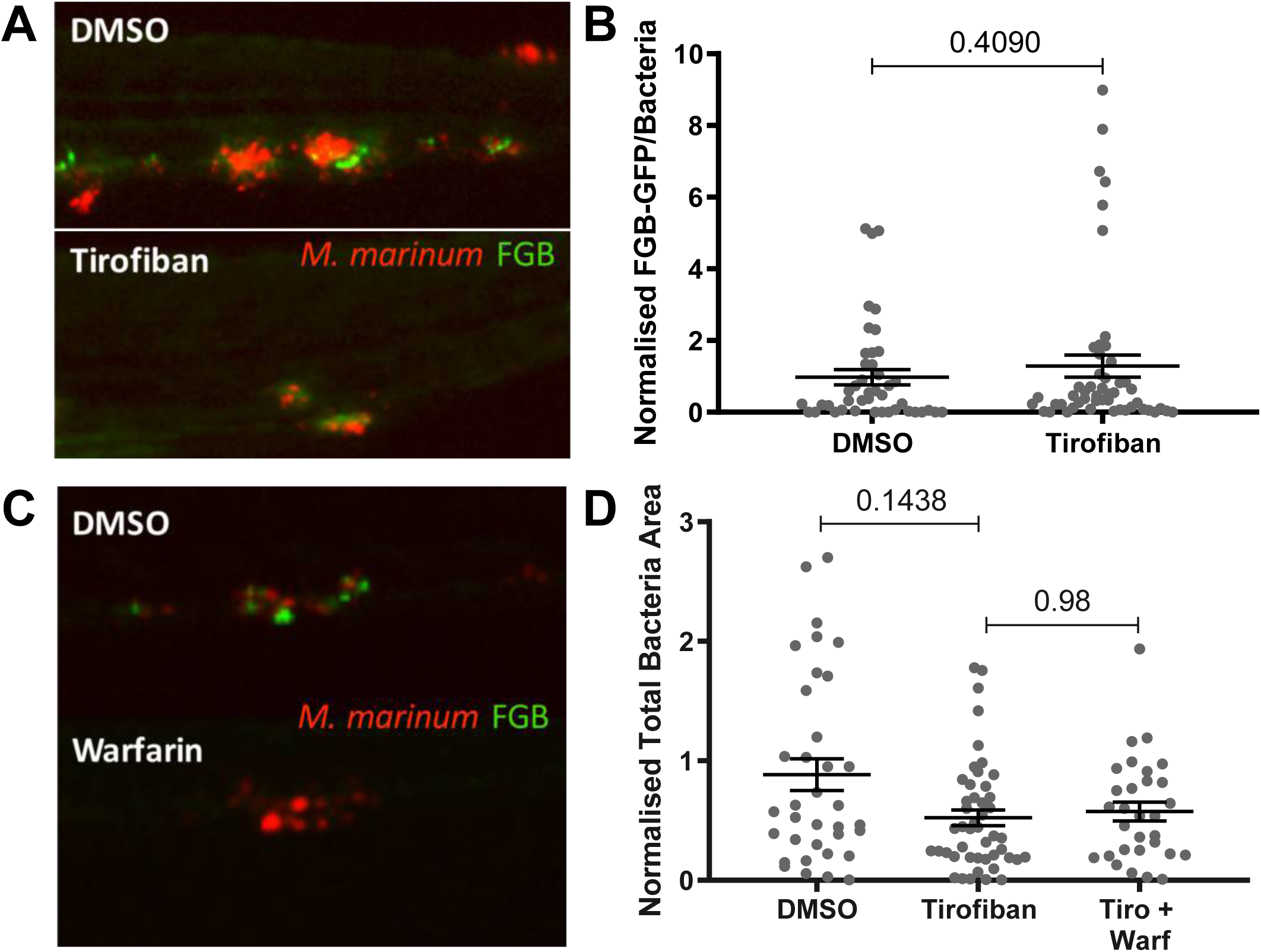
Thrombocytes increase mycobacterial burden independently of coagulation. A) Representative images of 5 DPI *Tg*(*fabp10a:fgb-EGFP)* embryos infected with *M*.*marinum*-tdTomato and treated with either DMSO or tirofiban. B) Quantification of clotting relative to bacterial burden in embryos treated with tirofiban normalized to DMSO control. Data are combined results of two independent experiments. C) Representative images of *Tg*(*fabp10a:fgb-EGFP)* embryos, where clot formation can be visualized by GFP fluorescence, infected with *M*. *marinum*-tdTomato (red) and treated with either DMSO or warfarin. D) Quantification of bacterial burden in embryos treated with tirofiban, warfarin, or tirofiban and warfarin, normalized to DMSO control. Data are combined results of 2 independent experiments. All graphs show mean ± SEM, statistical analysis by T test or ANOVA where appropriate.

### Thrombocytes compromise immunity through physical interactions with granuloma macrophages

To investigate the effect of glycoprotein IIb/IIIa inhibitors on thrombocyte-granuloma interactions, we first measured the effect of tirofiban on the infection-induced thrombocytosis phenotype. Surprisingly, tirofiban treatment increased thrombocytosis compared to the untreated infected group (Figure 6A). Increased thrombocyte density was also observed in infections with ΔESX1 mutant *M*. *marinum*, that are unable to drive granuloma maturation or necrosis, (Figure 6B), yet tirofiban treatment did not reduce ΔESX1 *M*. *marinum* burden (Figure 6C), further suggesting that the infection-induced thrombocytosis phenotype is independent of granuloma immunity.

**Figure 6:**
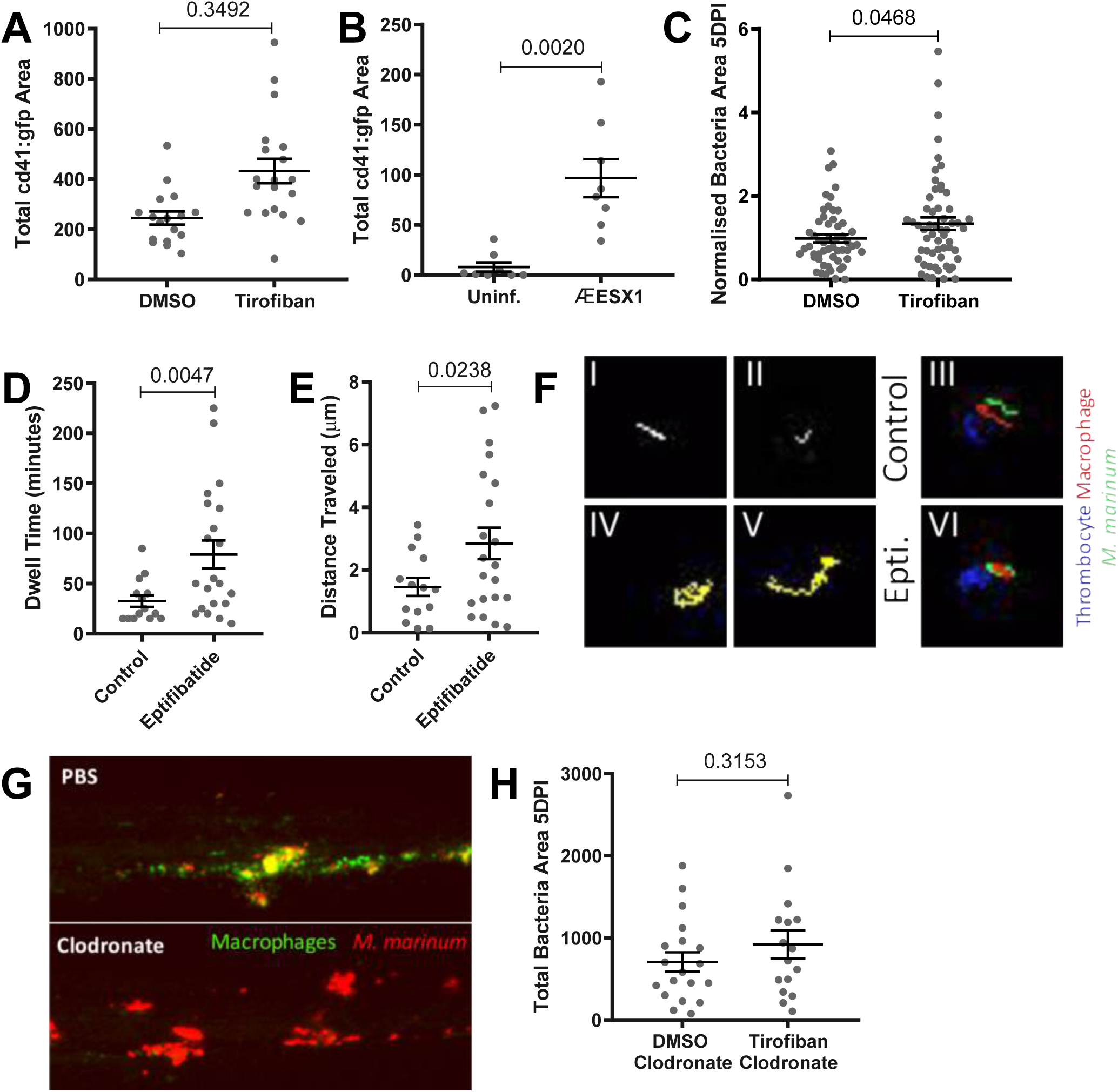
Thrombocyte activation compromises immunity through granuloma-associated macrophages. A) Quantification of total thrombocyte *Tg*(*cd41:GFP)* area within the tail WT *M*. *marinum*-tdTomato infected embryos treated with either DMSO or tirofiban. B) Quantification of total thrombocyte *Tg*(*cd41:GFP)* area within the tail of uninfected or ΔESX-1 *M*. *marinum*-tdTomato infected embryos. C) Quantification of bacterial burden in embryos infected with ΔESX1 *M*. *marinum*-tdTomato and treated with tirofiban. Combined data of 3 independent experiments. D) Quantification of length of thrombocyte association with an infected macrophage 5 DPI. Eptifibatide treatment was started at 4 DPI and was continued during timelapse. Data points represent individual thrombocytes from four (two control and two drug treated) infected fish. E) Quantification of distance thrombocyte traveled while associated with a site of macrophage and bacterial accumulation. Data points represent individual thrombocytes from four (two control and two drug treated) infected fish. F) Representative tracks of thrombocytes (I,II,IV,V) within control and eptifibatide treated embryos. Representative tracks of individual thrombocyte (blue), macrophage (red), and internalized *M*. *marinum* (green) (III, VI). G) Representative images of caudal hematopoietic tissue in 6 DPF *Tg*(*mfap4:tdTomato)* embryos, where macrophages are fluorescently labeled, injected with clodronate or PBS liposomes at 4 DPF. H) Quantification of bacterial burden in embryos infected with *M*. *marinum*-tdTomato, injected with clodronate liposomes and treated with tirofiban from at 3 DPI. All graphs show mean ± SEM, statistical tests by T-tests.

Given that glycoprotein IIb/IIIa-mediated platelet-monocyte binding can occur via fibrinogen ^23,24^ and patients with pulmonary tuberculosis have been shown to have significantly increased platelet-monocyte aggregation ^25^, we next aimed to determine whether anti-platelet drugs disrupted the interaction of thrombocytes and macrophages *in situ* within our model by live imaging. We infected double transgenic *Tg(cd41:GFP; mfap4:tdtomato)* with *M*. *marinum*-cerulean and performed timelapse imaging of 5 DPI embryos. Thrombocytes formed transient associations with *M*. *marinum*-infected macrophages (Supplementary Video 3), the frequency of which were increased upon eptifibatide treatment consistent with the increased thrombocytosis seen in Figure 6A (Supplementary Video 4), with an average dwell time of 30 minutes in untreated controls, and 80 minutes in eptifibatide-treated embryos (Figure 6D). We also observed a significant increase in the distance travelled by thrombocytes associated with *M*. *marinum*-infected macrophages in eptifibatide-treated embryos (Figure 6E). Thrombocyte tracks in control embryos were most often short and straight (Figure 6F I-II, Supplementary Video 5), maintaining a consistent distance from the center of the macrophage (Figure 6G III, Supplementary Video 6). In contrast, thrombocyte tracks in eptifibatide-treated embryos were long and circuitous, and varied in their distance from the center of the macrophage (Figure 6G IV-VI, Supplementary Videos 7 and 8).

To demonstrate that thrombocyte inhibition exerts a protective effect through boosting macrophage-dependent immunity, we depleted macrophages by injecting clodronate liposomes to deplete macrophages early during granuloma formation at 3 DPI (Figure 6C). Macrophage-depleted fish were unresponsive to tirofiban treatment demonstrating that pathological thrombocyte activation promotes bacterial growth via interactions with macrophages (Figure 6D). Together these results demonstrate glycoprotein IIb/IIIa inhibitor treatment disrupt pathological thrombocyte-macrophage attachments in mycobacterial granulomas.

### Granuloma maturation and pathological thrombocyte activation have a bidirectional relationship

Our observation that infection with ΔESX1 mutant *M*. *marinum* did not result in pathological thrombocyte activation (Figures 6C-D) suggested the existence of a bidirectional relationship between granuloma maturation, which is deficient in ΔESX1 mutant infections, and pathological thrombocyte activation. To further delineate this relationship we next took advantage of the stereotypical progression of innate immune granulomas in zebrafish embryos and used the burden reducing effect of tirofiban as a small molecule probe for thrombocyte activation. We found that at 3 DPI, a time-point with nascent granuloma formation but prior to significant granuloma organization and necrosis, tirofiban had no effect on bacterial burden (Figure 7A). Conversely, treatment of established infections from 4 to 5 DPI, a time-point when granulomas become organized and necrotic, tirofiban significantly reduced bacterial burden within 24 hours (Figure 7B). Together, these data demonstrate the existence of a switch point in granuloma maturity when thrombocytes are either activated or the activation of thrombocytes becomes pathological.

**Figure 7:**
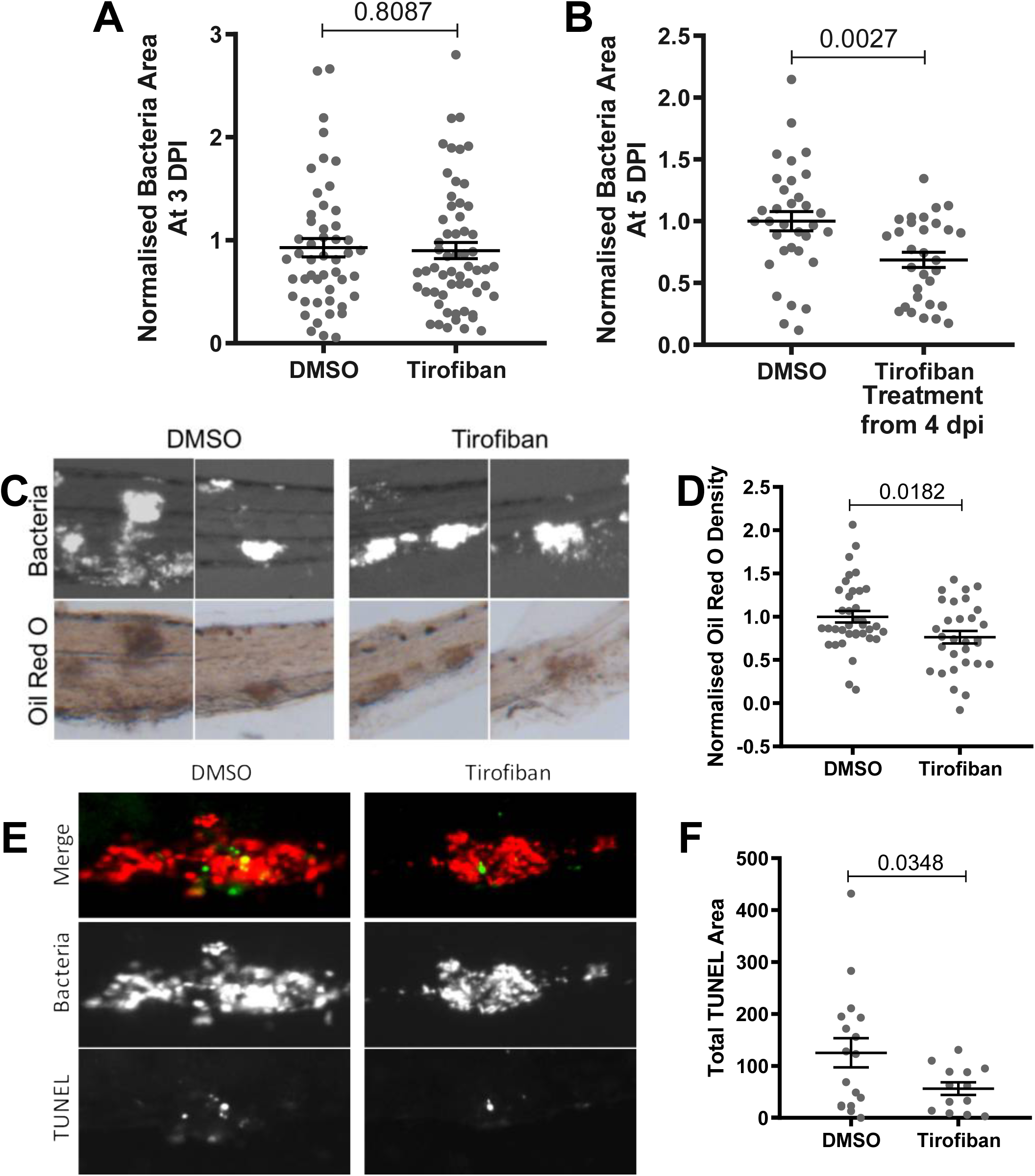
Granuloma maturation and pathological thrombocyte activation have a bidirectional relationship. A) Quantification of bacterial burden at 3 DPI after continuous tirofiban treatment from 0 DPI. B) Quantification of bacterial burden at 5 DPI after overnight drug treatment initiated at 4 DPI. C) Representative images of bacterial granulomas chosen for analysis (bacteria are white in greyscale images), and corresponding Oil Red O (ORO) staining (red-brown in color images). D) Quantification of ORO pixel density relative to granuloma bacterial area, in embryos treated with tirofiban, normalized to DMSO control. E) Representative images of bacterial granulomas showing bacteria in red and TUNEL staining in green. F) Quantification of TUNEL positive area within the largest granuloma of individual embryos. All graphs show Mean ± SEM, statistical tests by T-tests. Data are combined results of two independent experiments, except F) which represents a single experiment.

Co-incident with the appearance of granuloma necrosis at 4 to 5 DPI, we have demonstrated the appearance of foam cells in zebrafish embryo granulomas at this stage of infection ^15^. It has previously been shown that platelets accelerate the conversion of macrophages to foam cells in the presence of mycobacteria *in vitro* ^9^. Foam cells are permissive for mycobacterial growth, suggesting a mechanism for infection-induced thrombocyte activation to compromise innate immunity. We therefore hypothesized that thrombocyte inhibition would reduce the conversion of macrophages into foam cells. We investigated this by performing Oil-red O staining to measure lipid accumulation within size-matched granulomas (Figure 7C). Tirofiban-treated embryos had significantly less Oil-red O accumulation in their granulomas when compared to DMSO control, even after correction for reduced bacterial burden (Figure 7D). Together, these data demonstrate an *in vivo* effect of thrombocyte activation inhibiting an effective immune response by converting macrophages into foam cells in the maturing mycobacterial granuloma.

Given that foam cell formation is closely associated with necrosis in tuberculosis ^26^ we hypothesized that tirofiban treatment would reduce cell death within the granuloma. We therefore used TUNEL staining to detect the fragmented DNA of dying cells in *M*. *marinum* infected embryos. At 5 DPI, tirofiban-treated embryos showed significantly less TUNEL staining, indicating significantly reduced cell death within the granuloma (Figures 7E-F). Together these results indicate that infection-induced thrombocyte activation aggravates pathological markers of granuloma maturation and compromise immune control of mycobacterial infection.

## Discussion

Here we have used the zebrafish-*M*. *marinum* model to identify thrombocyte activation as a detrimental host response that is co-opted by pathogenic mycobacteria. Our data builds on previous studies that have shown coagulation, thrombocytosis and thrombocyte activation are associated with mycobacterial infection, and provides *in vivo* evidence of a direct role for thrombocyte activation in promoting mycobacterial growth. We have shown that infection-induced hemostasis is conserved in the zebrafish-*M*. *marinum* infection model, and that the platelet inhibiting drugs, aspirin, tirofiban, and eptifibatide, are able to reduce bacterial burden through host-mediated effects, independently of effects on coagulation.

A number of studies have investigated aspirin as a possible adjunctive treatment for TB in a range of animal models and human trials ^17-21,27,28^. The results of these studies have been far from conclusive, while most found beneficial effects ^17-19,21,27^, one human trial observed no effect ^20^, and a mouse study identified an antagonistic relationship between aspirin and the frontline anti-tubercular drug isoniazid ^28^. This lack of consensus may be due to the fact that the NSAID effect of aspirin will affect many cell types and processes important in the heterogeneous host response to mycobacterial infection. Our study expands this literature by delineating a role for glycoprotein IIb/IIIa in compromising the host response to infection.

Our study found that coagulation, thrombocytosis, and thrombocyte activation have distinct roles during the pathogenesis of mycobacterial infection of zebrafish. Inhibiting coagulation alone did not significantly reduce bacterial burden, and therefore we considered anti-platelet treatment as a more attractive HDT than anti-coagulant treatment. Although we found lower total clot formation in tirofiban-treated embryos, this was only proportional to bacterial load, suggesting infection-induced coagulation could be independent of infection-induced thrombocyte aggregation. It must be noted that we only measure a simple single end-point in our zebrafish embryo experiments (bacterial load) at a relatively early time point for a chronic infection. In more complex animals, where stroke and DVT are important secondary complications of mycobacterial infection, reducing coagulation may yet prove to be efficacious as a HDT during TB therapy to reduce morbidity. Conversely, data from the mouse model of TB suggests tissue factor-induced fibrin is necessary to contain mycobacteria within granulomas ^29^.

Our study provides evidence that while infection-induced thrombocytosis is a conserved function of infection, pathological thrombocyte-macrophage interactions are only driven in pathogenic mycobacteria during granuloma formation. Our experiments with ΔESX1 mutant *M*. *marinum*, which cannot secrete key virulence proteins that drive granuloma formation, demonstrated ESX1-dependent responsiveness to growth restriction by platelet inhibiting drugs. These data fit well with our observations that stationary thrombocytes were only observed around well-developed mycobacterial granulomas, and platelet inhibition was only effective at reducing bacterial burden after the development of significant granuloma pathology.

Our imaging and experimental data adds evidence that as for mammalian platelets, activated thrombocytes can form complexes with leukocytes through fibrinogen binding to glycoprotein IIb/IIIa, and that this alters immune cell function ^25,30-32^. Recent research has highlighted the important role of platelets as innate immune cells; they are able to release anti-microbial peptides, pick up and ‘bundle’ bacteria, and initiate the recruitment of other innate immune cells to sites of infection ^33-35^. However, the low frequency at which direct thrombocyte-mycobacterial interactions were observed in our study argues against thrombocytes having a significant role in directly mediating immunity to mycobacterial infection.

Platelets can induce macrophages to produce less pro-inflammatory TNF and IL-1ß, and more anti-inflammatory IL-10 in response to both BCG and *M*. *tuberculosis* ^7,9,25^. Crucially it has been shown that platelets are necessary for the formation of foam cells in the context of mycobacterial infection and atheroma ^9^. Our experiments demonstrating anti-platelet drug mediated control of infection, and reduced lipid accumulation and cell death in the granuloma suggest that infection-induced thrombocyte activation can be therapeutically modulated in mycobacterial infection.

Our findings that inhibition of thrombocyte activation reduces foam cell formation and cell death within the granuloma, leading to reduced bacterial burden provide important *in vivo* experimental evidence that infection-induced thrombocyte activation is a potential target for TB host-directed therapy. Specifically, our experimental demonstration of a thrombocyte-mediated effect of aspirin identifies a novel application for this well-tolerated and low cost drug in a disease of global importance.

## Supporting information

## Acknowledgements

We thank Dr Kristina Jahn and Sydney Cytometry for assistance with imaging equipment; Garvan Biological Testing Facility staff Ms Jennifer Brand, Mr Michael Pickering, Ms Rola Bazzi, Dr Lucie Nedved and Dr Stephanie Allison at the Garvan Institute of Medical Research for maintenance of zebrafish breeding stock; and Professors Lalita Ramakrishnan, Shaun Jackson and Georges Grau, Associate Professor Carl Feng, and Drs Jorn Coers and Imala Alwis for helpful discussion of results.

This work was supported by the Australian National Health and Medical Research Council APP1099912 and APP1053407; University of Sydney Fellowship; NSW Ministry of Health under the NSW Health Early-Mid Career Fellowships Scheme; and the Kenyon Family Inflammation Award (S.H.O.), Duke Summer Research Opportunities Program (K.J.), NIH Director’s New Innovator Award 1DP2-OD008614 (D.M.T), NIH R01-HL124232 and R01-HL125774 (J.A.S.).

## Author contributions

E.H., D.M.T. and S.H.O designed the experiments. E.H., K.E.J., M.D.J., T.N. and S.H.O performed the experiments. J.A.S. generated transgenic and mutant zebrafish lines. E.H. and S.O. wrote the paper. W.J.B., D.M.T. and S.H.O. supervised the project.

## Supplementary Figure Legends

**Figure S1.**
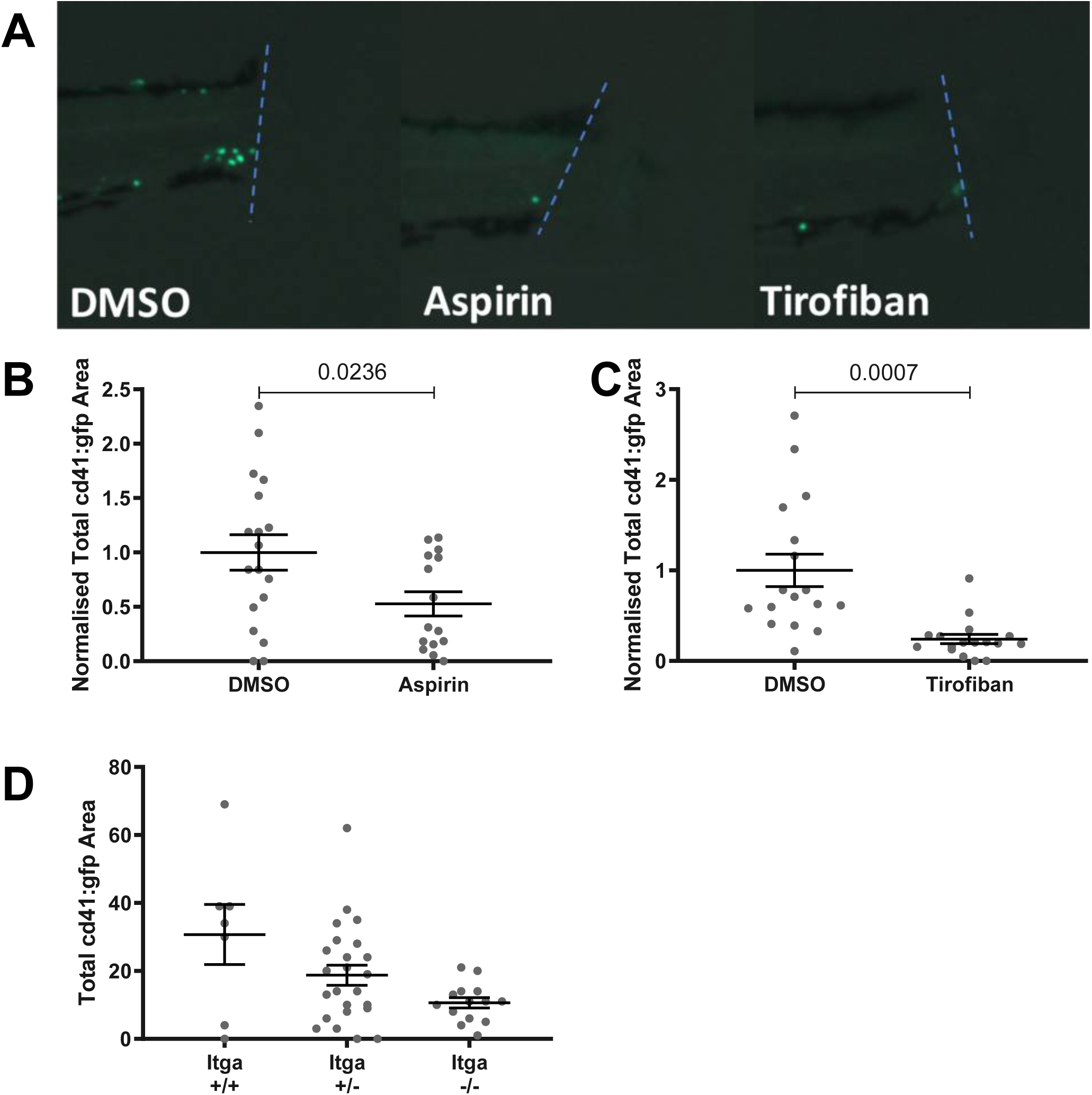
A) Representative images of thrombocyte accumulation at tail wound in 4 DPF *Tg*)*cd41:GFP)* embryos, 4 hours post injury (hpi). B) Quantification of thrombocytes within 100 μm of the cut site 4 hpi in aspirin-treated embryos. C) Quantification of thrombocytes within 100 μm of the cut site 4 hpi in tirofiban-treated embryos. D) Quantification of thrombocytes within 100 μm of the cut site 4 hours after injury in *itga2b* WT, heterozygous and knock-out embryos. Graphs show Mean ± SEM, statistical testing by T-test.

**Supplementary Video 1**

Video of 5 DPI *Tg*(*cd41:GFP*) embryo infected with *M*. *marinum*-tdtomato. Video captured with a long pass GFP filter to capture thrombocytes and bacteria simultaneously. Arrow indicates an example of a 5 minute thrombocyte-bacteria interaction.

**Supplementary Video 2**

Representative *Tg*(*fabp10a:fgb-EGFP)* embryo expressing EGFP-tagged fibrinogen beta (FGB) infected with *M*. *marinum*-tdtomato. Imaged every 15 minutes from 3 days post infection (DPI), until 6 DPI.

**Supplementary Video 3**

Representative timelapse of control *Tg*(*cd41:GFP; mfap4:tdtomato)* embryo infected with *M*. *marinum*-cerulean, showing the interaction of thrombocytes (green), macrophages (red), and bacteria (blue). Images were captured every 5 minutes for 4 hrs. Arrow shows successive short ‘binding’ events between thrombocytes and an infected macrophage.

**Supplementary Video 4**

Representative timelapse of eptifibatide treated *Tg*(*cd41:GFP; mfap4:tdtomato)* embryo infected with *M*. *marinum*-cerulean, showing the interaction of thrombocytes (green), macrophages (red), and bacteria (blue). Images were captured every 5 minutes for 5 hrs. Arrows show simultaneous long ‘binding’ events between thrombocytes and infected macrophages.

**Supplementary Video 5**

Representative tracks of thrombocytes associated with infected macrophages, from timelapse of control *Tg*(*cd41:GFP; mfap4:tdtomato)* embryo infected with *M*. *marinum*-cerulean. Images were captured every 5 minutes for 4 hrs, and tracks were generated using the Manual Tracking function in ImageJ.

**Supplementary Video 6**

Representative tracks of thrombocytes (blue and teal), showing the tracks of the macrophage (green and purple) and internalized M. marinum (red and yellow) they are associated with. From timelapse of control *Tg*(*cd41:GFP; mfap4:tdtomato)* embryo infected with *M*. *marinum*-cerulean. Images were captured every 5 minutes for 4 hrs, and tracks were generated using the Manual Tracking function in ImageJ.

**Supplementary Video 7**

Representative tracks of thrombocytes associated with infected macrophages, from timelapse of eptifibatide treated *Tg*(*cd41:GFP; mfap4:tdtomato)* embryo infected with *M*. *marinum*-cerulean. Images were captured every 5 minutes for 5 hrs, and tracks were generated using the Manual Tracking function in ImageJ.

**Supplementary Video 8**

Representative tracks of thrombocytes (blue and teal), showing the tracks of the macrophage (green and purple) and internalized M. marinum (red and yellow) they are associated with. From timelapse of eptifibatide treated *Tg*(*cd41:GFP; mfap4:tdtomato)* embryo infected with *M*. *marinum*-cerulean. Images were captured every 5 minutes for 4 hrs, and tracks were generated using the Manual Tracking function in ImageJ.

